# Differential regulation of TCR-induced ZFP36 and ZFP36L1 expression by cyclosporin A in CD8+ T cells

**DOI:** 10.1101/2025.09.24.678222

**Authors:** Marian Jones Evans, Twm J. Mitchell, Magdalena Zaucha, Georg Petkau, Martin Turner

## Abstract

CD8+ T cells target infected or malignant cells via the production of pro-inflammatory cytokines and direct target cell killing. Members of the ZFP36-family of RNA binding proteins, ZFP36 and ZFP36L1 regulate these functions in T cells via the regulation of mRNA stability and protein translation. We investigate the regulation of ZFP36 and ZFP36L1 expression using in vitro differentiated OT1 TCR transgenic memory-like T cells. We characterise the differential kinetics and sensitivity of ZFP36 and ZFP36L1 to antigen affinity and PMA versus ionomycin stimulation. By selectively inhibiting TCR-induced signalling pathways, we find that p38 MAPK, MEK1/2 and PKC contribute to inducing both ZFP36 and ZFP36L1 expression. By contrast, inhibition of calcineurin using cyclosporin A potently inhibits ZFP36L1 expression while increasing and prolonging ZFP36 expression. The Zfp36 promoter contains many binding sites for the transcription factors ELK-1/4 and few binding sites for NFAT, while the Zfp36l1 promoter contains many NFAT binding sites and few ELK1/4 binding sites. Our findings suggest that regulation of divergent transcription factors enable calcineurin to act as a signalling node that mediates the differential regulation of ZFP36 and ZFP36L1 during T cell activation.

## Introduction

T cell activation leads to proliferation and acquisition of effector functions which are essential for the elimination of infections and ultimately formation of memory. The T cell receptor is a key signal transducer which has been extensively characterised in terms of its proximal modes of signalling and downstream targets. One of the multiple pathways induced by TCR-antigen engagement includes the bifurcating pathway that hydrolyses phosphatidylinositol 4,5-bisphosphate into inositol trisphosphate and diacylglycerol. This leads to the activation of protein kinase C, the Ras-ERK Mitogen activated protein kinase (MAPK) pathways, and release of intracellular calcium (Ca2+) [1]. TCR stimulation also activates the p38 MAPK pathway and phosphoinositide 3-kinases (PI3K) which orchestrate numerous downstream events to drive the proliferation and differentiation of the activated T cell [1].

Early events after T cell activation include the nuclear translocation of transcription factors (TFs) NF-kB and NFAT that pre-exist in the cytoplasm in an inactive state. An additional class of targets regulated early after T cell activation are RNA Binding Proteins (RBPs) which profoundly influence gene expression in the immune system [2],[3]. The regulatory activity of TFs and RBPs drive a wave of rapid “immediate early” gene transcription and “delayed early” genes for which gene transcription is independent of *de novo* protein synthesis [4]. Together these pathways promote the transcription of secondary response genes - defined as genes which require *de novo* protein synthesis to be efficiently transcribed. Of TCR downstream targets, TFs and epigenetic regulators have been explored in the most detail, but some RBP are also rapidly responsive to TCR stimulation and these include ZFP36 which is an immediate early gene in non-lymphoid cells [5].

The ZFP36-family of RBP, three of which (ZFP36, ZfP36L1 and ZFP36L2) can be expressed by T cells, contain a tandem zinc finger RNA binding domain which interacts with AU-rich sequences in RNAs with high affinity to direct the localisation and translation or degradation of the bound RNA [2],[6]. These RBP have important roles in T cell function. They were initially shown to be repressors of cytokine production by T cells [7]–[12]. Subsequently, evidence has emerged that these RBP regulate many mRNAs in T cells and that these play a role in regulating the speed of T cell differentiation, the potency of cytotoxic T cells [13],[14] and the responsiveness of CD8 T cells to IL-2 [15].

ZFP36 and ZFP36L1 are rapidly and transiently induced following TCR engagement [13]–[15], unlike ZFP36L2, which is expressed in resting T cells and induced slowly following TCR stimulation [16]. The overlapping expression patterns of ZFP36 and ZFP36L1 are suggestive of functional redundancy and experiments using T cells lacking combinations of *Zfp36* and *Zfp36l1* have demonstrated redundancy [13],[14],[17],[18]. However, these and other studies [15],[19],[20], have also provided evidence for unique non-redundant roles, in particular in the context of differentiation and effector function. ZFP36L1 but not ZFP36 induction was shown to be highly sensitive to TCR affinity and critical for clonal selection of high affinity T cells [15]. How the expression of these RBPs is controlled by TCR signalling has not been investigated.

In addition to being transcribed from an immediate early gene, ZFP36 is also phosphorylated by MAP kinase-activated protein kinase 2 (MK2) downstream of RAS-MEK, and the p38 MAPK pathway [21],[22]. In a mouse macrophage cell line the p38 MAPK and ERK pathways synergistically induce ZFP36 protein, but not mRNA [23]. The phosphorylation of ZFP36 has been shown to promote its stability, while simultaneously inhibiting its ability to promote RNA decay. Regulation of ZFP36 by MAPK-induced phosphorylation has been shown to play a major role in modulating expression of *Tnf* and other pro-inflammatory mRNAs in macrophages [24]–[26]. MK2 also phosphorylates ZFP36L1 [27] as does the PI3K activated protein kinase B /AKT [28] and ERK-activated p90 ribosomal S6 kinase [29]. These post-translational events also promote the accumulation of ZFP36L1 protein [30]. It is unknown whether and how these pathways influence the expression of ZFP36 family members in activated T cells.

Here, using *in vitro* expanded antigen experienced CD8+ T cells we investigate the pathways that regulate ZFP36 and ZFP36L1 expression following TCR engagement. We identify calcineurin to be a point of divergent regulation between ZFP36 and ZFP36L1. We provide evidence that ZFP36L1 is principally an NFAT responsive gene, while ZFP36 is principally responsive to the ERK-ELK transcription factor axis. We suggest that downstream of the TCR, calcineurin switches off expression of the immediate early gene ZFP36 and induces the more sustained expression of ZFP36L1.

## Results

### TCR stimulation induces early and transient ZFP36, but sustained ZFP36L1 expression

Previous studies of ZFP36-family member expression in terminally differentiated effector CD8^+^ T cells showed that ZFP36 and ZFP36L1 proteins were both transiently induced after TCR stimulation within a 24-hour window [13]. To study this in non-terminally differentiated cells, we expanded naïve OT1 CD8^+^ T cells *in vitro* to generate large numbers of cells with a memory-like phenotype, as previously described [31] which could be used for flow cytometry, Western blotting and qPCR analysis following stimulation with peptide ligands which have a range of affinities for the OT1 TCR [32]. In these cells, TCR stimulation using the highest affinity N4 peptide caused rapid induction of *Zfp36* mRNA which peaked by one hour and decreased by two hours after stimulation (**Fig. 1A**). TCR agonists of lower affinity also rapidly induced *Zfp36* mRNA, but this remained higher at two hours than in cells stimulated with N4 peptide and declined thereafter. N4 only induced greater *Zfp36* mRNA than lower affinity antigens one hour after stimulation; at later timepoints *Zfp36* mRNA was not proportional to antigen affinity. In comparison, *Zfp36l1* mRNA was highest two hours after stimulation; at its peak the fold change in mRNA was greatest with the highest affinity peptide and decreased according to peptide affinity (N4 > T4 > Q4H7 > V4) (**Fig. 1B**). At later timepoints, fold change of the *Zfp36l1* mRNA induced by N4 continued to be greater compared to the lower affinity peptides. The kinetics of *Zfp36l1* mRNA expression were consistent irrespective of peptide affinity except for V4 which very weakly induced Zfp36l1. Thus, both transcripts are induced at the level of mRNA following TCR stimulation but with different kinetics and responses to antigen affinity.

**Figure 1.**
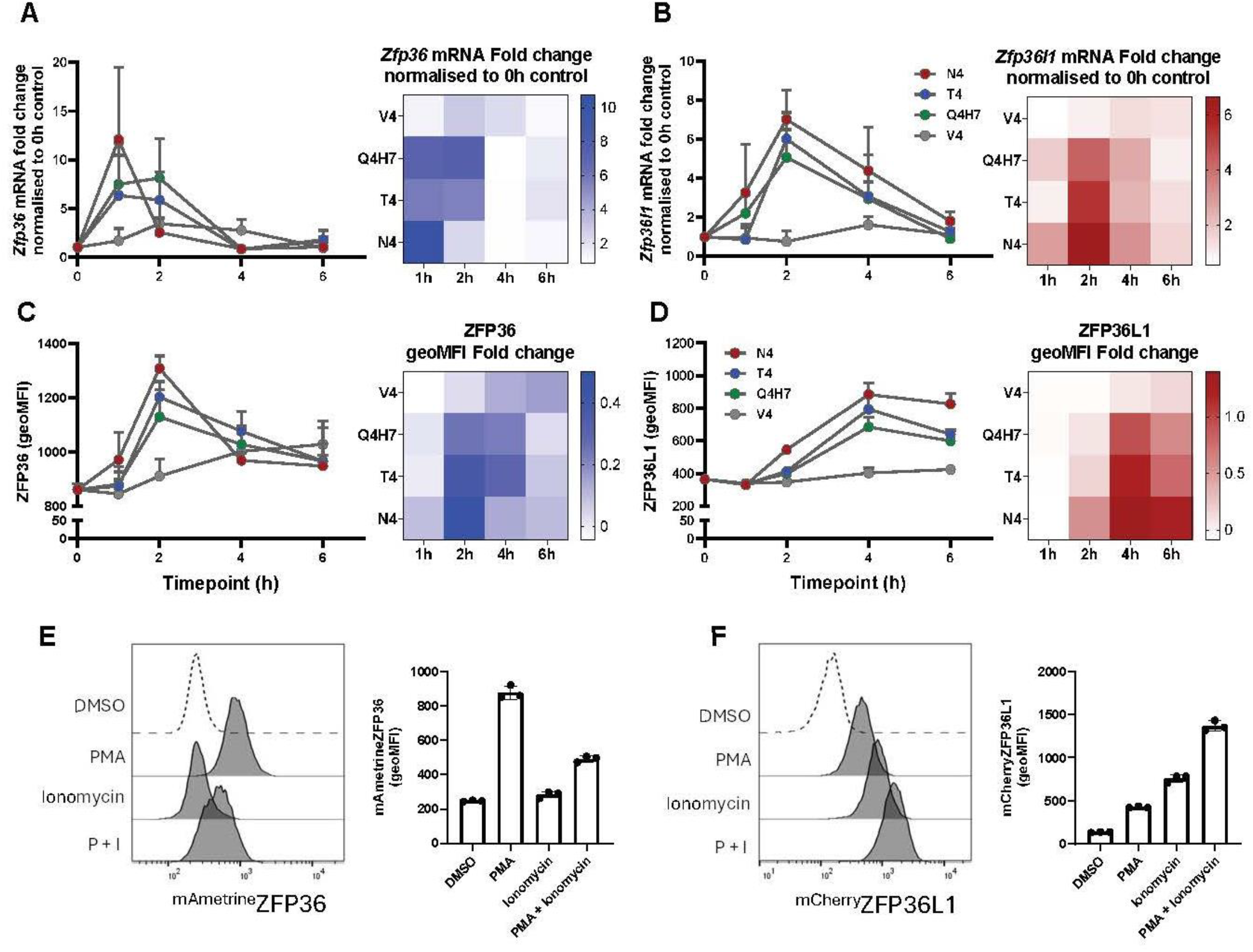
ZFP36 and ZFP36L1 expression exhibit distinct kinetics, sensitivity to antigen affinity and PKC *versus* Ca^2+^ signalling. Graphs show (A-B) *Zfp36* and *Zfp36l1* mRNA fold change, and (C-D) ZFP36 and ZFP36L1 geometric mean fluorescence intensity (geoMFI) in WT memory-like T cells stimulated for up to six hours with 0.1 nM N4, or lower affinity altered peptide ligands (T4, Q4H7, or V4), relative to unstimulated cells. mRNA fold change was quantified by the ΔΔCt qPCR method, and proteins were detected by intracellular flow cytometry. (A-D) Heat maps summarise fold change in (A-B) *Zfp36* and *Zfp36l1* mRNA or (C-D) ZFP36 and ZFP36L1 protein abundance. Error bars show standard deviation of the mean (SD) of three biological replicates. (E) Detection of ^mAmetrine^ZFP36 and (F) ^mCherry^ZFP36L1 in memory-like T cells stimulated for three hours with 10 ng/ml PMA and/or 1 µM ionomycin or treated with DMSO alone. Representative histograms show ^mAmetrine^ZFP36 or ^mCherry^ZFP36L1 expression from one of three homozygous biological replicates. The geoMFI was quantified and plotted; error bars show SD of three biological replicates.

We monitored the kinetics of protein expression at single cell resolution, in the same cells from which we harvested mRNA, by intracellular flow cytometry using antibodies specific for ZFP36 and ZFP36L1. The peak expression of ZFP36 was delayed compared to peak expression of its transcript (**Fig. 1C**). At this peak, but not thereafter, the amounts of ZFP36 scaled according to antigen affinity. ZFP36L1 also peaked later than its transcript and, at all timepoints analysed, the fold change of protein expression scaled according to peptide affinity (**Fig. 1D**). The differing kinetics of ZFP36 and ZFP36L1 expression were also reflected in the frequencies of ZFP36^+^ and ZFP36L1^+^ cells (**Fig.S1);** N4 stimulation induced the greatest proportion of cells expressing ZFP36 two hours post-stimulation. The frequency of ZFP36L1^+^ cells was consistently higher *versus* lower affinity peptides at all timepoints analysed. The data show that ZFP36 displays an expression pattern in memory-like cells that is consistent with it being an immediate early gene, while ZFP36L1 behaves as a delayed primary response gene [4] with expression linked to antigen affinity.

### ZFP36 and ZFP36L1 are distinctly sensitive to MAPK, PKC and Ca2+ signalling

To identify the mechanisms regulating ZFP36 and ZFP36L1 expression we used memory-like cells derived from genetically modified mice in which the *Zfp36* and *Zfp36l1* loci were modified by insertion of open reading frames encoding the fluorescent proteins mAmetrine or mCherry in the same reading frame as the start codon of ZFP36 or ZFP36L1 respectively [15]. These alleles retain the transcriptional and post-transcriptional regulatory elements of the endogenous transcripts, with the ^mAmetrine^ZFP36 and ^mCherry^ZFP36L1 reporter proteins predicted to be subject to the same post-translational regulation as the unmodified proteins. We compared reporter protein expression in memory-like T cells derived from ^mAmetrine^ZFP36 and ^mCherry^ZFP36L1 heterozygous or homozygous mice to validate that analysis of homozygous cells resulted in the greatest dynamic range of detection (**Fig. S2**). We therefore used homozygous mice for all analysis using ^mAmetrine^ZFP36 and ^mCherry^ZFP36L1 reporter proteins, unless stated otherwise.

We stimulated memory-like T cells with phorbol 12-myristate 13-acetate (PMA) and ionomycin which mimic the bifurcating signals of TCR signal transduction via Phospholipase C. PMA induces signalling via PKC and Ras pathways in T cells, while ionomycin leads to Ca^2+^ release and the activation of calcineurin. Three hours post-stimulation, PMA strongly induced ^mAmetrine^ZFP36 expression, while ionomycin alone stimulated very limited expression. Notably, ^mAmetrine^ZFP36 expression in response to combined PMA and ionomycin stimulation was less than that following stimulation with PMA alone (**Fig. 1E**). In contrast, ^mCherry^ZFP36L1 expression was more strongly induced by ionomycin alone than by PMA alone and these agents induced the greatest amounts of ^mCherry^ZFP36L1 when used together (**Fig. 1F**). We confirmed these observations at additional timepoints using Western blotting and found preferential induction of ZFP36 by PMA (**Fig. S3A**) and of ZFP36L1 by ionomycin (**Fig. S3B**) up to four hours post-stimulation. After one-hour, ZFP36 was strongly induced by PMA and decreased after two hours of stimulation. Ionomycin stimulation failed to induce ZFP36 expression that was detectable using our conditions for Western blotting. When added together with PMA, ionomycin reduced slightly ZFP36 expression (**Fig. S3A**). Thus, ZFP36 and ZFP36L1 respond differently to PMA and ionomycin.

We further investigated the signalling pathways regulating the expression of ZFP36 and ZFP36L1 in the presence of small molecule inhibitors which were added to memory-like T cells prior to TCR stimulation. Four hours after stimulation, BIRB 796 (p38 MAPK inhibitor), trametinib (MEK1/2 inhibitor) and Go6983 (PKC inhibitor) each inhibited expression of ^mAmetrine^ZFP36 (**Fig. 2A - C**). Inhibiting p38 MAPK or MEK1/2 pathways also inhibited ^mCherry^ZFP36L1 induction (**Fig. 2D - E**). Inhibition of PKC signaling resulted in a modest inhibition of ^mCherry^ZFP36L1 expression (**Fig. 2F**) compared to the effect of Go6983 on ^mAmetrine^ZFP36 expression. Taken together, these data indicate that ZFP36 and ZFP36L1 are both induced by p38 MAPK, MEK1/2 and PKC signalling. The inhibition of the upstream signalling node PKC dominantly regulates ZFP36 expression, but not ZFP36L1, suggesting non-redundancy of this pathway for the expression of ZFP36. These pathways are likely to underlie the ability of PMA to induce both ZFP36 and ZFP36L1.

**Figure 2.**
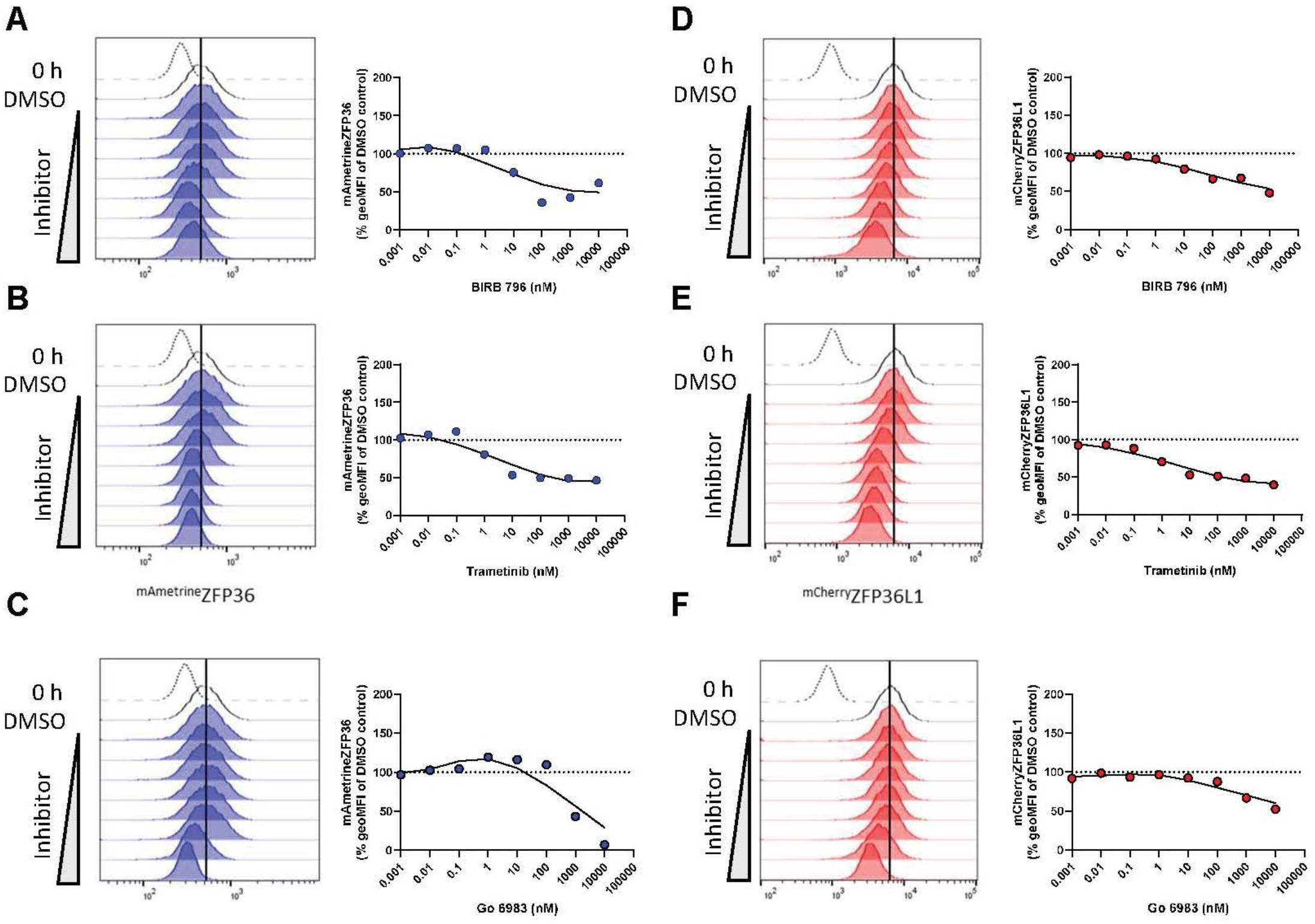
p38 MAPK, MEK1/2 and PKC signalling are common pathways that promote the expression of both ZFP36 and ZFP36L1. Memory-like T cells were pre-treated for 30 minutes with 0.001, 0.01, 0.1, 1, 10, 100, 1000, or 10,000 nM BIRB 796 (p38 MAPK inhibitor), Go6983 (PKC inhibitor) or trametinib (MEK1/2 inhibitor) or DMSO before being stimulated for four hours with 0.1 nM N4 in the presence of each inhibitor. Histograms show (**A - C**) ^mAmetrine^ZFP36 geoMFI or **(D – F)** ^mCherry^ZFP36L1 expression after four hours of stimulation. Broken line shows 0 h unstimulated control and unfilled histogram shows stimulated DMSO-only control. Filled histogram shows ^mAmetrine^ZFP36 or mCherryZFP36L1 geoMFI of cells stimulated in the presence of the indicated inhibitor. The effect of each inhibitor on ^mAmetrine^ZFP36 or ^mCherry^ZFP36L geoMFI was analysed by normalising the geoMFI following inhibitor treatment as a percentage of the DMSO-treated stimulated control. The ^mAmetrine^ZFP36 or ^mCherry^ZFP36L1 geoMFI of resting cells was subtracted as background fluorescence prior to calculating the percentage inhibition of expression. Data shown is from one biological replicate and is representative of a total of three ^mAmetrine^ZFP36 ^mCherry^ZFP36L1 homozygous biological replicates.

### Calcineurin differentially regulates ZFP36 and ZFP36L1 expression

While PMA stimulation alone was sufficient to induce both ZFP36 and ZFP36L1 expression (**Fig. 1E-F**), ZFP36 and ZFP36L1 exhibited contrasting sensitivity to ionomycin. We therefore further investigated the role of TCR-induced Ca^2+^ signalling in regulating ZFP36 and ZFP36L1 expression through use of the calcineurin inhibitor cyclosporin A (CsA). Strikingly, ZFP36 expression four hours post-stimulation was higher in the presence of CsA than in cells treated with DMSO (**Fig. 3A**). By contrast, CsA was a potent dose-dependent inhibitor of ^mCherry^ZFP36L1 induction by TCR stimulation (**Fig. 3B**). CsA also increased ^mAmetrine^ZFP36 expression at six- and eight-hours after stimulation; this effect was increasingly pronounced as, in the absence of CsA, ^mAmetrine^ZFP36 expression was declining (**Fig. 3C**). As observed at four-hours, the inhibition of ZFP36L1 expression by CsA was consistent at six- and eight-hours post-stimulation (**Fig. 3D**). Thus, CsA inhibits ^mCherry^ZFP36L1 expression but sustains the expression of ZFP36.

**Figure 3.**
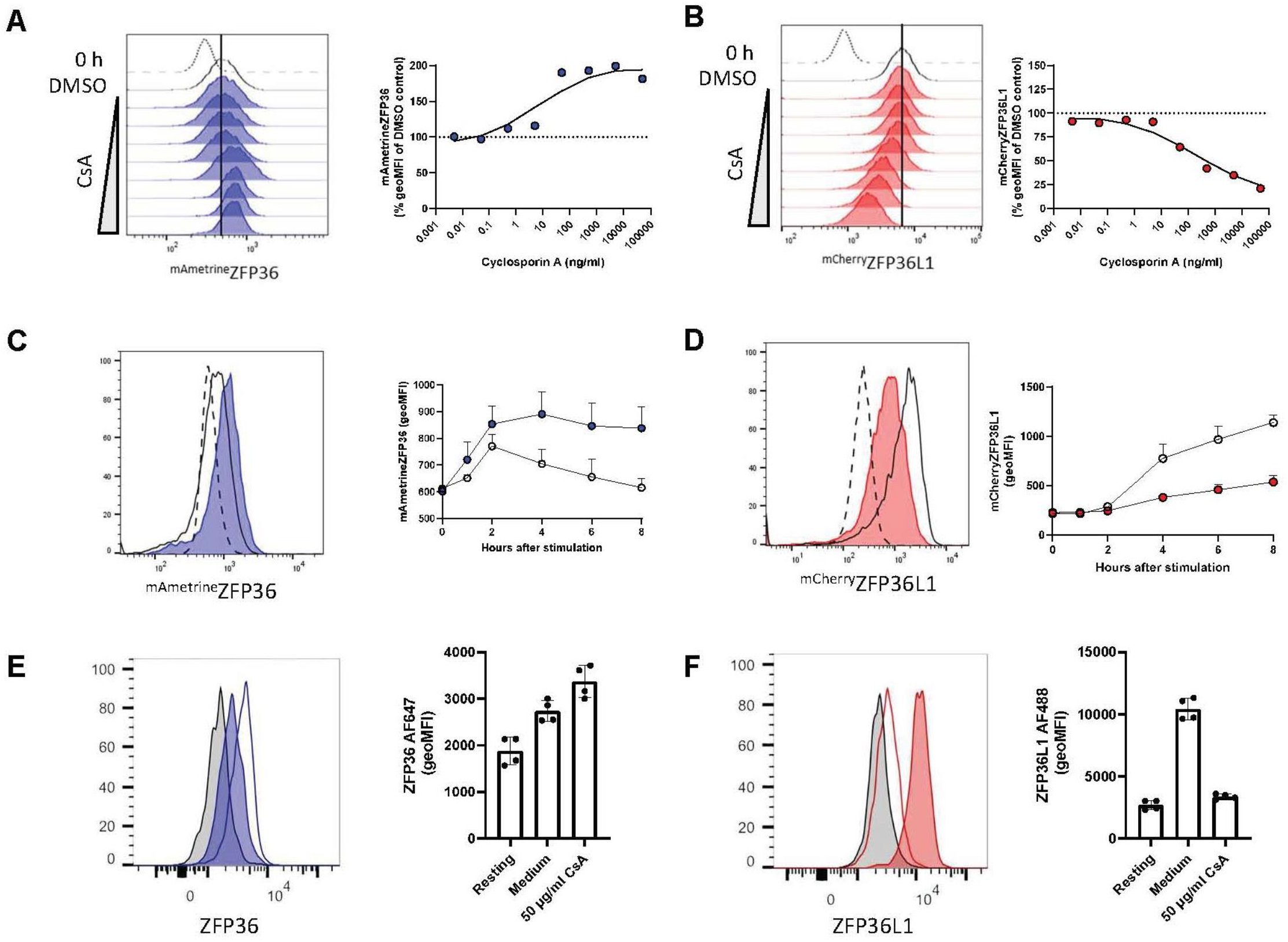
Cyclosporin A increases ZFP36 and inhibits ZFP36L1 expression. **(A – B)** Memory-like T cells were pre-treated for 30 minutes with 0.005, 0.05 0.5, 5, 50, 500, 5000 or 50,000 ng/ml cyclosporin A (CsA) (calcineurin inhibitor) or DMSO control before being stimulated for four hours with 0.1 nM N4 in the presence of CsA or DMSO. Histograms show ^mAmetrine^ZFP36 or ^mCherry^ZFP36L1 geoMFI after four hours of stimulation. Broken line shows 0-hour unstimulated control and unfilled histogram shows stimulated DMSO control. Filled histogram shows geoMFI of cells stimulated in the presence of CsA. For quantification, the effect of CsA was analysed by normalising the ^mAmetrine^ZFP36 or ^mCherry^ZFP36L1 geoMFI following inhibitor treatment as a percentage of the DMSO-treated stimulated control. The ^mAmetrine^ZFP36 or ^mCherry^ZFP36L1 geoMFI of resting cells was subtracted as background fluorescence prior to calculating the percentage inhibition of expression. Data shown is from one biological replicate and is representative of a total of three homozygous biological replicates. (**C - D**) Memory-like T cells were pre-treated for 30 minutes with 500 ng/ml CsA before being stimulated for up to 8 hours with 0.1 nM N4 in the presence of CsA or DMSO. Representative histograms show ^mAmetrine^ZFP36 and ^mCherry^ZFP36L1 expression from one of three homozygous biological replicates. Broken line shows 0-hour unstimulated control and unfilled histogram shows stimulated DMSO control. Filled histogram shows geoMFI of cells stimulated in the presence of CsA. The geoMFI was quantified and plotted; error bars show standard deviation of the mean (SD) of three biological replicates. (**E – F**) *Ex vivo* memory T cells were stimulated for three hours with 10 ng/ml PMA and 1 µM ionomycin in the presence or absence of CsA. The CsA-treated cells were pre-treated with CsA for 30-minutes prior to stimulation. ZFP36 and ZFP36L1 expression in *ex vivo* memory T cells was analysed by flow cytometry analysis by pre-gating on CD8^+^ CD44^high^ CD62L^low^ to identify the population of memory T cells. Representative histograms show (**E**) ZFP36 or (**F**) ZFP36L1 expression in unstimulated (light grey), or PMA/ionomycin stimulated memory cells in the absence (filled coloured histogram) and presence (open histogram) of CsA in one of four biological replicates. ZFP36 and ZFP36L1 geoMFI was quantified and the geoMFI of the four biological replicates plotted. Error bars show standard deviation of the mean (SD).

To extend our observations, we analyzed memory-like cells from mice expressing ^mAmetrine^ZFP36 and ^mCherry^ZFP36L1 by Western blotting in parallel with flow cytometry. N4 stimulation induced expression of ^mAmetrine^ZFP36 two hours post-stimulation which declined and was undetectable after eight hours (**Fig. S3C**). In the presence of CsA, ^mAmetrine^ZFP36 expression was greater after two hours of stimulation compared to untreated cells. ^mAmetrine^ZFP36 expression was also prolonged by the presence of CsA as increased ^mAmetrine^ZFP36 expression was maintained up to eight hours post-stimulation (**Fig. S3C**). CsA inhibited N4-induced ^mCherry^ZFP36L1 expression most potently two – four hours after stimulation and notably reduced the accumulation of protein resolving at a lower molecular weight (**Fig. S3C**) which may represent hypo-phosphorylated ^mCherry^ZFP36L1.

To confirm the effects of CsA on the endogenous ZFP36 and ZFP36L1 proteins we stimulated memory-like cells derived from wildtype OTI mice *in vitro*. CsA inhibited ZFP36L1 and prolonged the expression of ZFP36 (**Fig. S3D**). We also investigated if CsA similarly regulated ZFP36 and ZFP36L1 expression in *ex vivo* memory T cells. CsA treatment increased ZFP36 expression (**Fig. 3E**) while potently inhibiting ZFP36L1 expression to levels similar to that of resting cells (**Fig. 3F**). The effects of CsA suggest that Ca^2+^ signaling via calcineurin phosphatase is a key pathway that mediates the differential regulation of ZFP36 and ZFP36L1.

### Calcineurin regulates the expression of Zfp36 and Zfp36l1 mRNAs

Both the rate of transcription and transcript stability determine the overall abundance of mRNA. To gain mechanistic understanding of how CsA differentially regulates ZFP36 and ZFP36L1 expression, we investigated the effect of CsA on *Zfp36* and *Zfp36l1* mRNA abundance using qPCR analysis. The expression of *Zfp36* and *Zfp36l1* in resting cells was unaffected by CsA **(Fig. 4A)**. Following peptide stimulation, CsA treatment extended the duration of *Zfp36* mRNA expression which remained elevated at four- and six-hours post-stimulation without increasing the peak abundance two hours after stimulation. By contrast, CsA limited the induction of *Zfp36l1* mRNA to a maximum of 4-fold *versus* 14-fold in DMSO treated control cells (**Fig. 4B**). ZFP36 protein expression is subject to strong post-transcriptional regulation which includes its own autoregulation [33]. Deletion of AU-rich elements in the 3’UTR of *Zfp36* resulted in increased ZFP36 protein and greater resilience of mice to inflammation-induced disease [34]. The protein synthesis inhibitor cycloheximide (CHX) is thought to enhance *Zfp36* mRNA expression by inhibiting a negative feedback mechanism, mediated by *de novo* translated proteins including ZFP36 itself, that promotes *Zfp36* mRNA decay (**Fig. S5) [**35] [33]. CHX should abolish the effect of CsA if upon T cell activation CsA acts as an inhibitor of these negative regulators of *Zfp36* mRNA. To test whether negative feedback regulators of *Zfp36* mRNA are affected by CsA, we pre-treated memory like cells with CHX. The abundance of both *Zfp36* and *Zfp36l*1 mRNA was increased upon pretreatment with CHX between 2 and 6h after stimulation. However, the addition of CsA in the presence of CHX further increased the expression of *Zfp36* mRNA but not *Zfp36l1* mRNA (**Fig. 4C**). This suggests that induction of *Zfp36* mRNA expression by CsA is not via the inhibition of a negative feedback mechanism mediated by ZFP36 or other regulators of mRNA induced upon T cell activation.

**Figure 4.**
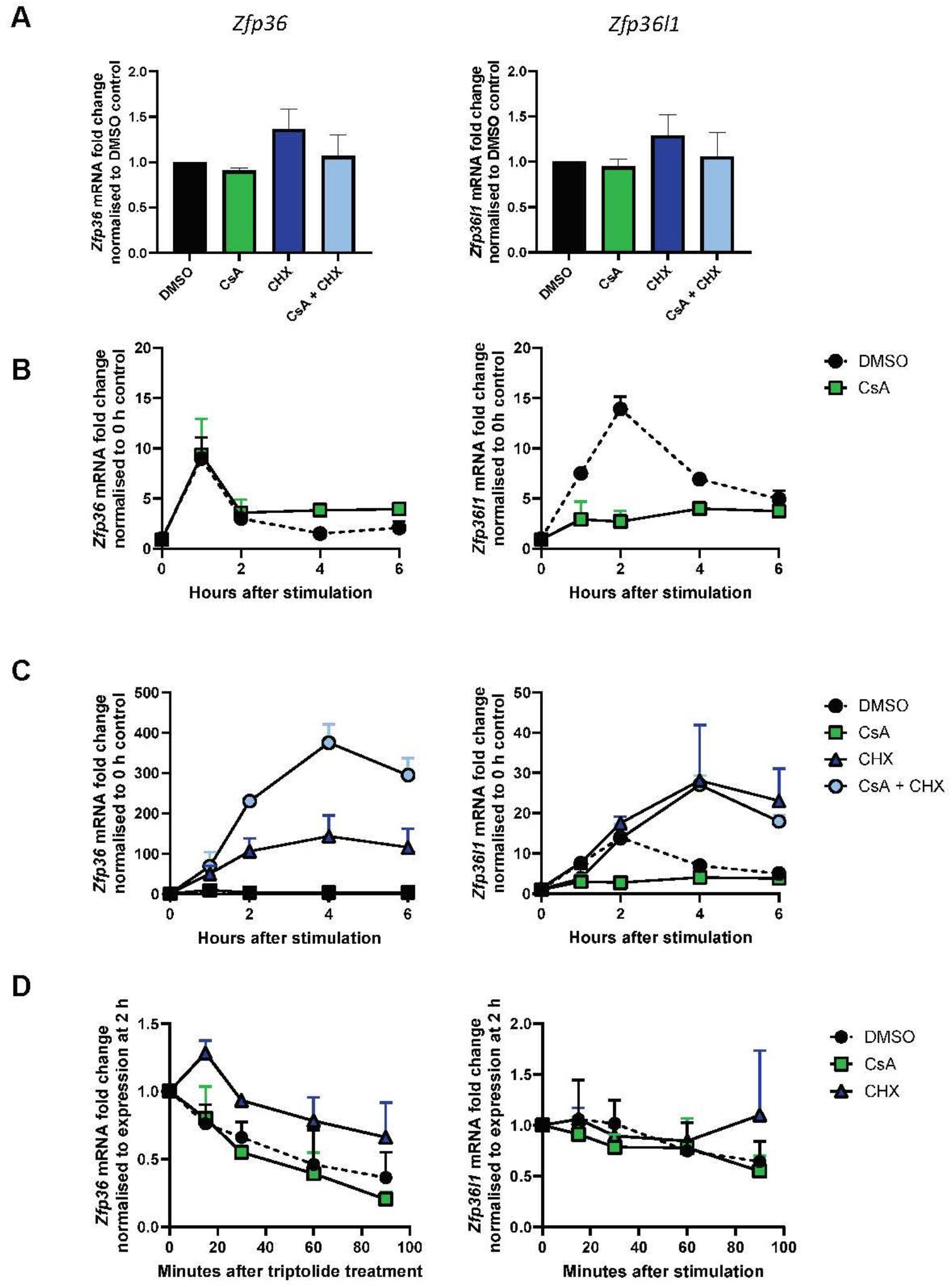
CsA prolongs *Zfp36* mRNA expression independently of de novo protein translation or transcript stability. (**A**) Abundance of *Zfp36* and *Zfp36l1* mRNA was measured in resting WT OTI memory-like T cells that were pre-treated for 30 minutes with 500 ng/ml cyclosporin A (CsA) (calcineurin inhibitor), 10 µg/ml cycloheximide (CHX) (translation inhibitor) or DMSO control, analysed by ΔΔCt qPCR analysis. (**B - D**) WT OTI memory-like T cells were stimulated for up to 6 hours with 0.1 nM N4 in the presence of 500 ng/ml CsA, 10 µg/ml CHX or DMSO control as indicated. (**B - C**) Abundance of *Zfp36* and *Zfp36l1* mRNA was determined by ΔΔCt qPCR analysis using cells pre-treated with DMSO as a universal control. (**D**) After 2 hours of stimulation, the cells were treated with 1 µM triptolide (transcription inhibitor) for up to 90 minutes. Fold-change of *Zfp36* and *Zfp36l1* mRNA abundance was normalised to the abundance in each condition 2 hours after stimulation, prior to triptolide treatment using ΔΔCt qPCR analysis. (**A - D**) Error bars show standard deviation of the mean of three biological replicates.

We further investigated if CsA modulates ZFP36 expression *via* stabilization of *Zfp36* mRNA. Inhibition of transcription using triptolide (TRP), which causes rapid proteasomal degradation of RNA polymerase II, enables analysis of mRNA stability [36]. We analysed the stability of *Zfp36* or *Zfp36l1* mRNA between two- and four-hours after stimulation, when the *Zfp36* transcript declines in DMSO-treated control cells but not in the presence of CsA (**Fig. 4B**). 90 minutes after TRP treatment, the abundance of *Zfp36* mRNA had rapidly declined 64 % in the DMSO-treated control compared to 80 % in the presence of CsA. In contrast, higher *Zfp36* mRNA abundance was maintained for longer following CHX treatment which increased the half-life of *Zfp36* mRNA to be greater than 90 minutes. Thus, CHX but not CsA treatment resulted in increased *Zfp36* mRNA stability. The stability *Zfp36l1* mRNA was not affected by the presence of either CsA or CHX (**Fig. 4D**). We therefore conclude that the induction of ZFP36 expression by CsA is not due to the post-transcriptional stabilisation of the *Zfp36* transcript.

ZFP36 protein stability can be regulated by MAPK-dependent and -independent pathways [37],[38]. We therefore investigated if the inhibition of calcineurin promotes the accumulation of ZFP36 protein. To analyse the effect of CsA on the stability of ZFP36 protein and its degradation, we used either cycloheximide or the proteasome inhibitor MG-132 to inhibit *de novo* translation or block degradation of ZFP36 respectively. We added these compounds after two hours of stimulation with N4, when ^mAmetrine^ZFP36 expression is near its maximum. The addition of CHX had limited effect on ^mAmetrine^ZFP36 compared to the DMSO control at two – six hours after stimulation, indicating that the majority of ZFP36 protein has already been translated in the first two-hours of stimulation and that it is relatively stable (**Fig. S6A-B)**. MG-132 treatment increased ^mAmetrine^ZFP36 expression at four – six hours post-stimulation, indicating that ZFP36 is subject to proteasomal degradation at these timepoints (**Fig. S6C)**. These data indicate there is little impact of CsA on ZFP36 protein stability. We conclude that the effects CsA are mostly at the level of *Zfp36* mRNA transcription.

### Calcineurin sensitive transcription factors can explain the differential expression of ZFP36 and ZFP36L1 in response to CsA

Next, we investigated if the opposing regulation of *Zfp36* and *Zfp36*l1 mRNA may be mediated by different transcription factors which regulate *Zfp36* and *Zfp36l1*. As MEK1/2 and p38 MAPK are key pathways regulating ZFP36 expression in memory-like T cells (**Fig. 2A - B**) we considered the transcription factor ELK-1, a member of the ternary complex factors (TCFs) subfamily of transcription factors, which has been shown to promote *Zfp36* mRNA expression in MCF-7 cells downstream of ERK1/2 activation in response to epidermal growth factor (EGF) stimulation [39]. In double positive thymocytes lacking ELK-1 and ELK-4 *Zfp36* expression was 0.5-fold lower as a ratio compared to WT following plate-bound anti-CD3 stimulation [40] (**Fig. 5A**). *Zfp36l1* mRNA showed only minor reduction in the absence of ELK-1 and ELK-4. Moreover, there was no additive effect on *Zfp36l1* mRNA in double deficient thymocytes (**Fig. 5B**). Promoter analysis using the Eukaryotic Promoter Database (Dreos et al., 2014) identified three putative binding sites for both ELK-1 and ELK-4 that were within a few hundred bases of the *Zfp36* transcription start site and thus could promote *Zfp36* transcription (**Fig. 5C**). The *Zfp36l1* promoter region contains multiple ELK-4 binding sites but just two ELK-1 binding sites, one of which lies just downstream of the transcription start site (**Fig. 5D**). The greater number and position of the ELK-1 binding sites in the *Zfp36* promoter compared to the *Zfp36l1* promoter suggests that ELK-1 may have a greater role in regulating *Zfp36* than *Zfp36l1* transcription. Notably, two independent studies demonstrated that ELK-1 is dephosphorylated by calcineurin at phosphoserine 383 (S383) and that this inhibits its transcriptional activity [41],[42]. We therefore propose that the prolonged expression of *Zfp36* mRNA in the presence of CsA is, at least in part, due to loss of calcineurin-mediated ELK-1 inhibition.

**Figure 5.**
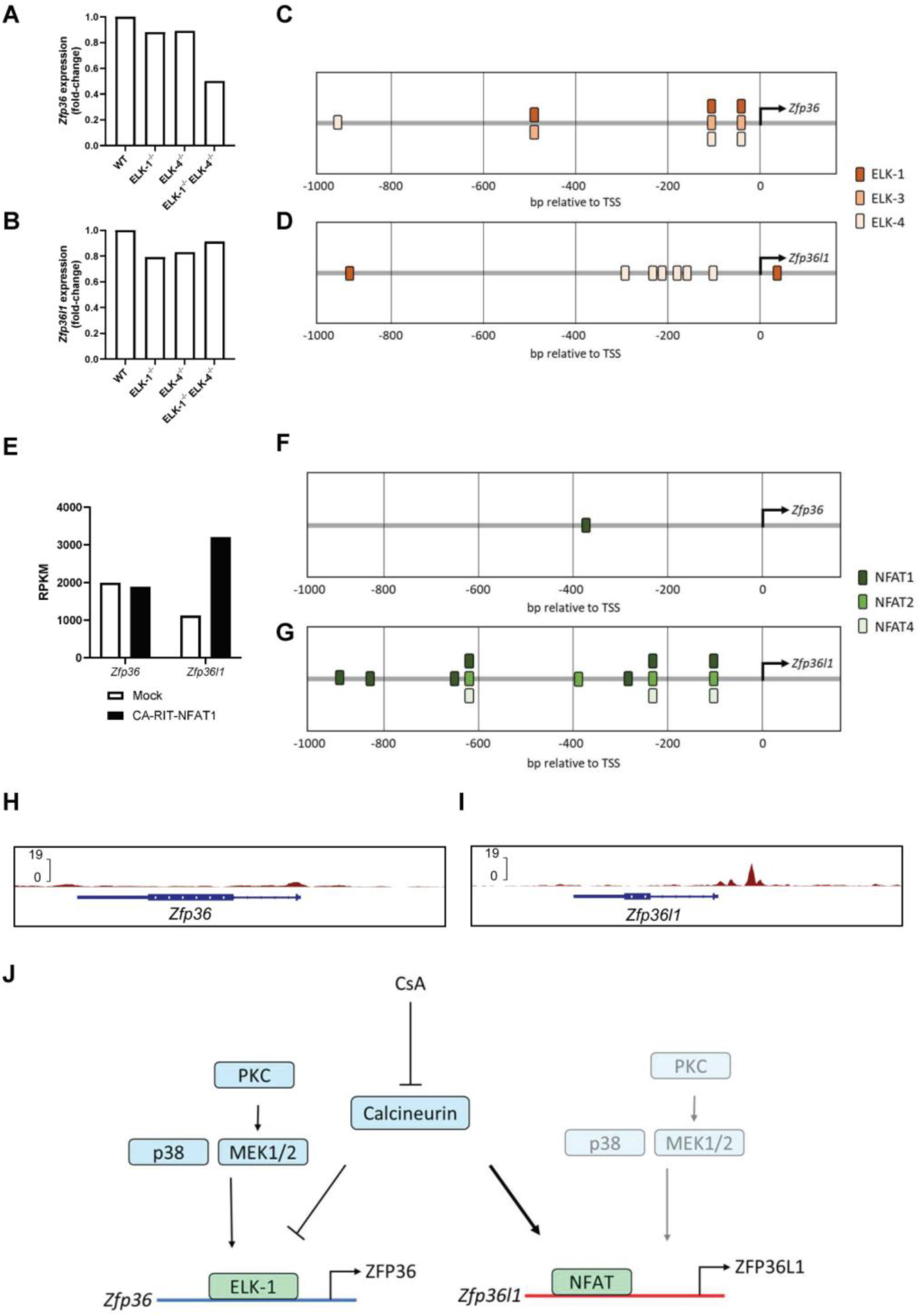
ELK and NFAT may underlie divergent regulation of *Zfp36* and *Zfp36l1* by calcineurin. (**A**) *Zfp36* or (**B**) *Zfp36l1* microarray probe signal induced by 30-minute plate-bound CD3 stimulation in ELK mutant double positive thymocytes as a ratio compared to wildtype (WT). Data from Costello et al., 2010. (**C - D**) Graphical representation of ELK-family transcription factor binding sites in the (**C**) *Zfp36* and (**D**) *Zfp36l1* promoters; analysis from the Eukaryotic Promoter Database[53] using a cut-off p-value of 0.001. (**E**) RPKM values from RNA-Seq for *Zfp36* and *Zfp36l1* expression in CA-RIT-NFAT1-transduced or mock-transduced CD8^+^ T cells taken from ref-[45] (**F - G**) Graphical representation of NFAT-family transcription factor binding sites in the (**F**)*Zfp36* and (**G**) *Zfp36l1* promoters as in panel (**C - D**). (**H - I**) ChIP-seq data from ref-[45] for endogenous NFAT binding to (**H**) *Zfp36* or (**I**) *Zfp36l1* in WT cells stimulated with PMA and ionomycin for one hour. (**J**) Visualisation of the proposed mechanisms underlying the negative regulation of ZFP36 and induction of ZFP36L1 by calcineurin. Activation of ELK-1 by p38 MAPK, MEK1/2 and PKC signalling promotes the transcription of *Zfp36*. Calcineurin negatively regulates ELK-1 to inhibit *Zfp36* transcription. In contrast, calcineurin promotes NFAT-mediated *Zfp36l1* transcription and is the dominant pathway inducing ZFP36 expression compared to a lesser role of p38 MAPK, MEK1/2 and PKC. Inhibition of calcineurin by CsA results in increased ZFP36 and inhibits ZFP36L1 expression.

Dephosphorylation of NFAT-family transcription factors by calcineurin facilitates their nuclear translocation and transcriptional activity [43]. When NFAT interacts with Fos-Jun (AP-1), cooperative NFAT:AP-1 complexes are formed which have been shown to promote the induction of cytokines such as IL-2 [44],[45]. To examine the effect of NFAT on *Zfp36* and *Zfp36l1* we examined RNA-seq data generated from CD8^+^ T cells expressing a constitutively active form of NFAT1 which cannot interact with AP-1 [45]. In these cells the constitutively active NFAT1 promotes increased expression of *Zfp36l1*, but *Zfp36* expression is unchanged (**Fig. 5E**). Analysis of the promoter regions of *Zfp36 and Zfp36l1* identifies only a single NFAT1 binding site in the *Zfp36* gene (**Fig. 5F**) compared to seven NFAT1, four NFAT2 and three NFAT4 binding sites in the *Zfp36l1* gene (**Fig. 5G**)). Furthermore, CHIP-seq data for NFAT in T cells [45] shows much greater NFAT binding to the *Zfp36l1* gene promoter than to the *Zfp36* gene (**Fig. 5H - I**). The enrichment of NFAT binding sites in the *Zfp36l1* compared to *Zfp36* gene is consistent with *Zfp36l1* being more responsive to ionomycin than *Zfp36*.

## Discussion

Here, we provide evidence that calcineurin regulates ZFP36 and ZFP36L1 expression in CD8^+^ memory-like T cells by enforcing the transient expression of ZFP36 while promoting sustained ZFP36L1 expression. Taken together, our data lead us to propose a model for the divergent regulation of ZFP36 and ZFP36L1 expression mediated by the calcineurin-responsive ELK-family and NFAT-family transcription factors (**Fig. 5J**). This provisional model will require additional validation focussing on the regulation of transcription factors and the transcription of the *Zfp36* and *Zfp36l1* genes in activated T cells and exploration of whether calcineurin inhibitors can prolong ZFP36 expression in non-T cells.

Given that CsA is a potent immunosuppressant [46] and that both ZFP36 and ZFP36L1 have previously been shown to limit T cell mediated inflammatory and immune responses in a redundant manner [13],[14],[17],[18], our observations are relevant to the mechanism of action of CsA as an immunosuppressant. The immunosuppressive mechanism of action of CsA is partly explained by the inhibition of NFAT-mediated transcription required for T cell activation [43]. However, CsA was also reported to inhibit the cytokines TNF-β and lymphotoxin-β independently of an effect on NFAT [47]. By inhibiting the dephosphorylation of ELK1, CsA prolongs the expression of *Zfp36* mRNA enabling the protein to be expressed for an extended period. Interestingly, CsA has also been reported to inhibit the activation of p38 MAPK [48],[49]. In this scenario, when p38/MK2 mediated phosphorylation of ZFP36 is diminished the RBP is better able to promote mRNA decay. Thus, CsA may extend the expression and function of ZFP36 beyond its narrow duration as an immediate early gene to limit T cell activation. This suggestion is consistent with the observation that mice with prolonged ZFP36 expression, resulting from removal of the autoinhibitory sequence in the *Zfp36* 3’UTR show much reduced disease severity in and experimental autoimmune encephalitis a T cell-driven autoimmune disease [34].

Expression of ZFP36L1 in response to ionomycin stimulation alone, and in response to the constitutively active, AP-1 independent, NFAT mutant indicates that NFAT activation alone is sufficient for ZFP36L1 expression. The extent to which ZFP36L1 is part of the programme of T cell anergy/tolerance or exhaustion is interesting to consider as ZFP36L1 plays a role in promoting the response to IL-2 and limiting effector cell differentiation [15] and posttranscriptional silencing of cytokine mRNA has been linked to the phenotype of anergic cells [50], but the underlying mechanisms remain uncharacterized.

In summary, the distinct regulation of ZFP36 and ZFP36L1 early after T cell activation suggest they play unique roles in shaping the dynamic transcriptome of the activated T cell.

## Materials and Methods

### Mice

Mice with modified Zfp36 and Zfp36l1 alleles acting as protein expression reporters have been previously described: Zfp36tm2.1Tnr (MGI:7711587); (MGI:7325470) [15], with the OT-I TCR-transgene (Vα2 and Vβ5 recognizing peptide residues 257-264 of chicken ovalbumin in the context of H-2Kb) (MGI:3054907) and CD4cre transgene (MGI:2386448) were maintained on the C57BL/6 background at the Babraham Institute.

All mouse experimentation was approved by the Babraham Institute Animal Welfare and Ethical Review Body. Animal husbandry and experimentation complied with existing European Union and United Kingdom Home Office legislation and local standards under project licence PP9973990). Mice were bred and maintained in the Babraham Institute Biological Support Unit. Since the opening of this barrier facility (2009), no primary pathogens or additional agents listed in the FELASA recommendations have been confirmed during health monitoring surveys of the stock holding rooms. Ambient temperature was ~19-21°C and relative humidity 52%. Lighting was provided on a 12-hour light: 12-hour dark cycle including 15 min ‘dawn’ and ‘dusk’ periods of subdued lighting. After weaning, mice were transferred to individually ventilated cages with 1-5 mice per cage. Mice were fed CRM (P) VP diet (Special Diet Services) ad libitum and received seeds (e.g. sunflower, millet) at the time of cage-cleaning as part of their environmental enrichment. Male and female mice were used in the experiments described here; all mice were aged between 6- and 20-weeks of age.

### Antibodies and Flow Cytometry

For cell surface staining single cell suspensions from cultured cells were prepared in FACS buffer (DPBS + 2 % FBS + 2 mM EDTA). All cells were blocked with Fc blocking antibody (24G2, BioXcell) and incubated with fixable cell viability dye eF780 (Thermo Fisher or BD) for 30 minutes at 4 °C.

For intracellular staining of ZFP36 and ZFP36L1, cells were surface stained before being fixed with Cytofix/Cytoperm Fixation/Permeabilization Kit (BD) for 30 minutes at 4°C. Intracellular staining was performed in Permwash (BD) containing the intracellular antibody cocktail overnight at 4 °C. The following antibody clones were used in the flow cytometry experiments: CD8 (53-6.7), CD44 (IM7), CD69 (H1.2F3), CD62L (MEL-14) ZFP36 (D1I3T), ZFP36L1 (E6L6S). Data was acquired using a ZE5 Cell Analyzer (Bio-Rad) or a Fortessa (Beckton Dickinson) flow cytometer equipped with 355, 405, 488, 561, and 640 nm lasers. Flow Cytometry data was analysed using FlowJo 10.8.1 software.

### In vitro culture and stimulation of memory-like T cells

Single cell suspensions of cells from the spleen and lymph nodes (inguinal, axillary and brachial) of mice were prepared by mechanical disruption of harvested tissues against a 40 μM cell strainer. Tissue harvesting and preparation of single cell suspensions were performed using RPMI-1640 medium (Thermo Scientific) supplemented with 2 % FBS (Thermo Scientific) and 2 mM EDTA (Thermo Scientific). Selective lysis of red blood cells using ACK lysis buffer (Thermo Scientific) was performed to enrich for leukocytes.

The engineered SAMBOK (also referred to as MEC.B7.SigOVA) mouse embryonic fibroblast (MEF) cell line which expresses OVA_257-264_ presented by H-2K^b^ and the co-stimulatory ligand CD80 were used as APCs for the generation of memory-like T cells as previously described (van Stipdonk et al., 2003). SAMBOK cells were maintained in RPMI-1640 media (Thermo Scientific) supplemented with 10 % FBS and passaged by 1:15 dilution twice per week from a confluent density of 10 × 10^6^ cells/T75 flask by trypsinisation (Gibco TrypLE) for 5 minutes at 37 °C.

The day prior to T cell activation (Day (D).−1), 0.5 × 10^6^ SAMBOK cells were seeded per well in a 12-well plate. The following day (D.0), the SAMBOK cells were washed twice with PBS (Sarstedt) to remove loosely adhered cells. Isolated leukocytes were seeded at a density of 2 × 10^6^ cells/ml in a total volume of 2 ml per well of a 12-well plate. Cultures were maintained in Iscove’s Modified Dulbecco’s Medium (IMDM) (Thermo Scientific) medium supplemented with 10% FBS, 50 μM 2-mercaptoethanol (Thermo Scientific), 100 U/ml penicillin, and 100 μg/ml streptomycin (Thermo Scientific). The cells were cultured at 37°C, 5% CO_2_. After 24 hours (D.1), the cells in each well were gently resuspended and removed from the SAMBOK feeder layer before being combined in a 50 ml conical tube. The cells were centrifuged at 300 g for 5 minutes at room temperature. The cells were seeded at a density of 1 × 10^6^ cells/ml in IMDM media containing 10 ng/ml recombinant murine IL-7 (PeproTech or STEMCELL) in a 24-well plate containing 2 ml/well. Three days post-activation (D.3), the cells were reseeded in fresh IL-7 containing IMDM media at 1 × 10^6^ cells/ml in a final volume of 2 ml. From the fourth day of culture (D.4), 1 ml of the supernatant was discarded and replaced with 1 ml of fresh IMDM containing a 2x concentration of 20 ng/ml IL-7. All *in vitro* assays were performed on memory-like T cells between days 7 – 10 of culture. For all cell counting, cell number, diameter and percentage viability were quantified using a CASY Model TT cell counter (Scharfe systems/Roche). 10 μl of cell suspension was added to 10 ml of CASY ton. Cells between 7.5 – 13 µm in diameter were measured.

Memory-like CD8+ T cells were harvested and washed with pre-warmed supplemented IMDM and counted as previously described. For analysis of by flow cytometry, cells were seeded at a final density of 1 × 10^6^ cells/ml in 96-well U-bottom plates. For western blot or qPCR analyses, cells were seeded at a final density of 1 × 10^6^ − 4 × 10^6^ cells/ml in 24-well plates. For all assays, IL-7 was used at a final concentration of 10 ng/ml. For analysis of antigen responses, memory-like T cells were stimulated with N4 OVA peptide (Genscript) or the lower affinity variant OVA peptides T4, Q4H7, or V4. Alternatively, memory-like T cells were stimulated with PMA or ionomycin. Stimulation mixes were prepared as 2 × solutions which were diluted 1:2 upon addition to the plated cells.

Inhibitor dilutions were first prepared as 1000 × stocks by performing eight-point 1:10 serial dilutions of each inhibitor in DMSO (Sigma). For preparation of 2 × inhibitor solutions, a 1:500 further dilution was performed of each inhibitor in IMDM media. This method maintained a constant concentration of DMSO across the range of inhibitor concentrations. To facilitate the necessary series of dilutions, cells were initially plated at 4 × the final desired density with 40 ng/ml IL-7 and rested for 30 minutes at 37°C and 5% CO_2_. The cells were pre-treated for 30 minutes with a 2 × stock of the relevant inhibitor or DMSO. Activation solutions containing a 1 × inhibitor concentration and 2 × N4 concentration were prepared by combining equal volumes of the appropriate 2 × inhibitor solution (or DMSO) and 4 × N4 IMDM solution (or IMDM media alone). Equal volume of the activation solutions were added to the pre-treated cells in culture and returned to the incubator. The inhibitors used were BIRB 796 (Insight Biotechnology); CsA (Sigma-Aldrich); Go 698 (inhibits PKCα, PKCβ, PKCγ, PKCδ and PKCζ) (Insight Biotechnology); and Trametinib (Insight Biotechnology).

### Analysis of the effect of CsA on ex vivo memory T cells

Splenocytes were isolated from 20-week-old WT mice according to the method described above to generate single-cell suspensions. The splenocytes were pre-treated with 50 µg/ml CsA for 30 minutes prior to stimulation with 10 ng/ml PMA and 1 µM ionomycin. For flow cytometry analysis of ZFP36 and ZFP36L1, CD8^+^ memory T cells were gated according to the strategy described in **Figure S4**. The experiment was performed with two biological replicates, with two technical replicates respectively.

### Inhibition of mRNA transcription, protein translation or proteasomal degradation

To analyse mRNA stability, memory-like T cells were pre-treated with 500 ng/ml CsA or DMSO for 30 minutes. The cells were stimulated for 2 hours with N4 peptide before 1 µM triptolide (CST) or DMSO was added at a final concentration of for up to 90 minutes. To analyse superinduction of *Zfp36* and *Zfp36l1* mRNA, memory-like T cells were pre-treated for 30 minutes with 500 ng/ml CsA, 10 µg/ml cycloheximide (CHX) (Sigma-Aldrich) or DMSO. The cells were then stimulated with N4 for up to 6 hours. The cells were then harvested and frozen on dry ice for qPCR analysis.

To inhibit translation and enable analysis of protein stability, the cells were first pre-treated for 30 minutes with 500 ng/ml CsA or DMSO. Memory-like T cells were stimulated for 2 hours before 10 µg/ml CHX or DMSO was added, and the cells stimulated up to a further 4 hours. To inhibit proteasomal degradation, memory-like T cells were stimulated for 2 hours prior to addition of 10 μM MG-132 (Sigma-Aldrich) or DMSO and the cells were stimulated for up to a further 4 hours. The cells were harvested and prepared for flow cytometry analysis.

### Western Blotting

Whole cell lysates were prepared using 1x RIPA buffer [SDS 0.1 % (w/v), Sodium deoxycholate 0.4% (w/v), NP-40 0.5% (v/v), 100mM NaCl, 50mM Tris-HCl, pH 7.4], supplemented with 1:100 protease inhibitor cocktail (Sigma), phosphatase inhibitor cocktail III (Sigma), and 2 U/ml Benzonase (Sigma). Protein lysate concentration was determined using Pierce BCA protein assay kit (Thermo Scientific). Lysates were denatured for 5 min at 97°C with 4 × Laemmli buffer (200 mM Tris-HCl (pH 6.8), 8 % w/v SDS, 40 % v/v glycerol, 0.08 % w/v bromophenol blue) containing 5 % β2-mercaptoethanol. Protein lysate concentrations were normalized to enable the loading of an equivalent mass of protein per lane of a 10 % polyacrylamide SDS-PAGE gel. 5 μl of precision plus protein dual colour standard (Bio-Rad) was loaded in one lane as a molecular weight marker. 1 μg of protein lysates of HEK293 cells expressing FLAG-tagged mouse ZFP36 or ZFP36L1 was loaded as a positive control for antibody staining. The proteins were resolved by running at 120 V for ~1.5 hours before being transferred to a nitrocellulose membrane using the iBlot II transfer device (Thermo Scientific) at 15 V for 6 min. Ponceau staining was performed to visualise protein transfer before the membrane was washed with Milli-Q water. The membrane was incubated in intercept TBS blocking buffer (Li-Cor), a ready-to-use blocker formulation, for 2 hours at room temperature with constant shaking. Following blocking, the membrane was incubated with primary antibodies diluted in intercept buffer overnight at 4°C with constant shaking. The membrane was washed three times with TBS-T (0.05% Tween-20), and three times with TBS, before being incubated with secondary antibodies diluted in intercept buffer for 2 hours at room temperature. The membrane was washed again before image acquisition using the Odyssey CLx (Li-Cor) and analysed using the ImageStudio Lite version 5.2 (Li-Cor).

The membranes were probed using the following primary antibodies: ZFP36 (Origene 3D10), ZFP36L1 (CST BRF1/2 or E6L6S), GAPDH (CST D16H11) and α-tubulin (CST DM1A)). Secondary antibody staining was performed using the following antibodies:, anti-mouse IgG IRDye800CM (Licor) and anti-rabbit IgG IRDye680RD (Licor).

### Real Time PCR analysis

For generation of samples for qPCR analysis, cells were thoroughly resuspended and harvested into at least twice the volume ice-cold PBS in 15 ml Falcon tubes. The cells were centrifuged at 300 g for 5 minutes at 4 °C before the supernatant was discarded. The cells were washed in PBS and transferred to a 1.5 ml Eppendorf tube before being centrifuged at 13,000 g for 1 minute at 4 °C. The supernatant was removed and the cell pellets snap-frozen on dry ice and stored at – 70 °C. RNA was extracted using the RNeasy Mini Kit (Qiagen) according to the manufacturer’s instructions including optional on-column DNAse treatment using the RNase-Free DNase Set (Qiagen). RNA concentration was quantified using NanoDrop One (Nanodrop). RNA concentrations were normalised prior to cDNA synthesis using the RT RevertAid first strand cDNA synthesis kit (Thermo Scientific) according to the manufacturer’s instructions. cDNA synthesis reactions were performed using the T100 Thermal Cycler (Bio-Rad) using the following programme: 25 °C for 5 minutes, 42 °C for 60 minutes and 70 °C for 5 minutes. Quantitative real-time PCR was performed using TaqMan universal PCR master mix (Thermo Scientific) and probes detailed below. Reactions were performed as technical triplicates in 384-well plates with nuclease free water (Qiagen) used as a no template control (NTC). qPCR reactions were performed using the CFX384 Touch Real-Time PCR Detection System (Bio-Rad). The thermal cycling parameters were as follows: 50 °C for 2 minutes, 95 °C for 10 minutes, 40 cycles of 95 °C for 15 seconds followed by 60 °C for 1 minute. The data was exported using the CFX Manager Software v. 3.1 (Bio-Rad) and analysed in Excel v.2408 (Microsoft) where the data was analysed according to the ΔΔCt method using *Rpl32 (*Mm07306626_gH) as a housekeeping gene to normalise against. qPCR primers for *Zfp36 (*Mm00457144_m1) and *Zfp36l1*

## Supporting information

supplementary figures combined

## Author Contributions and Notes

M.J.E. conceptualization; methodology; investigation; validation; formal analysis; visualisation; writing original draft preparation; John Evans et al., 23/09/2025 – preprint copy – BioRxiv T.J.M. conceptualization; methodology; G.P. conceptualization and writing, M.Z. conceptualization; investigation; formal analysis; M.T. conceptualization, supervision, funding acquisition, writing review and editing.

## Acknowledgments

We thank Sarah Bell, Michael Screen, Alexander Saveliev and Louise Matheson for their help in completing this project, the Babraham Institute Biological Support Unit and Flow Cytometry facilities for their outstanding support, Patrick Costello and David Bending for discussion and advice. Gemma Cassettari formatted this manuscript version for BioRxiv, using a template generously provided by the Finkelstein Lab, Austin, TX. This study was supported by funding from the Biotechnology and Biological Sciences Research Council (BBSRC) BB/P013414/1 to M.T.; the BBSRC Core Capability Grant to the Babraham Institute; and a Wellcome Discovery grant (226660/Z/22/Z) to M.T. M.J.E. was supported by a Babraham Institute studentship as part of the Cambridge BBSRC doctoral training programme.

