## supplementary figures combined for "Differential regulation of TCR-induced ZFP36 and ZFP36L1 expression by cyclosporin A in CD8+ T cells"

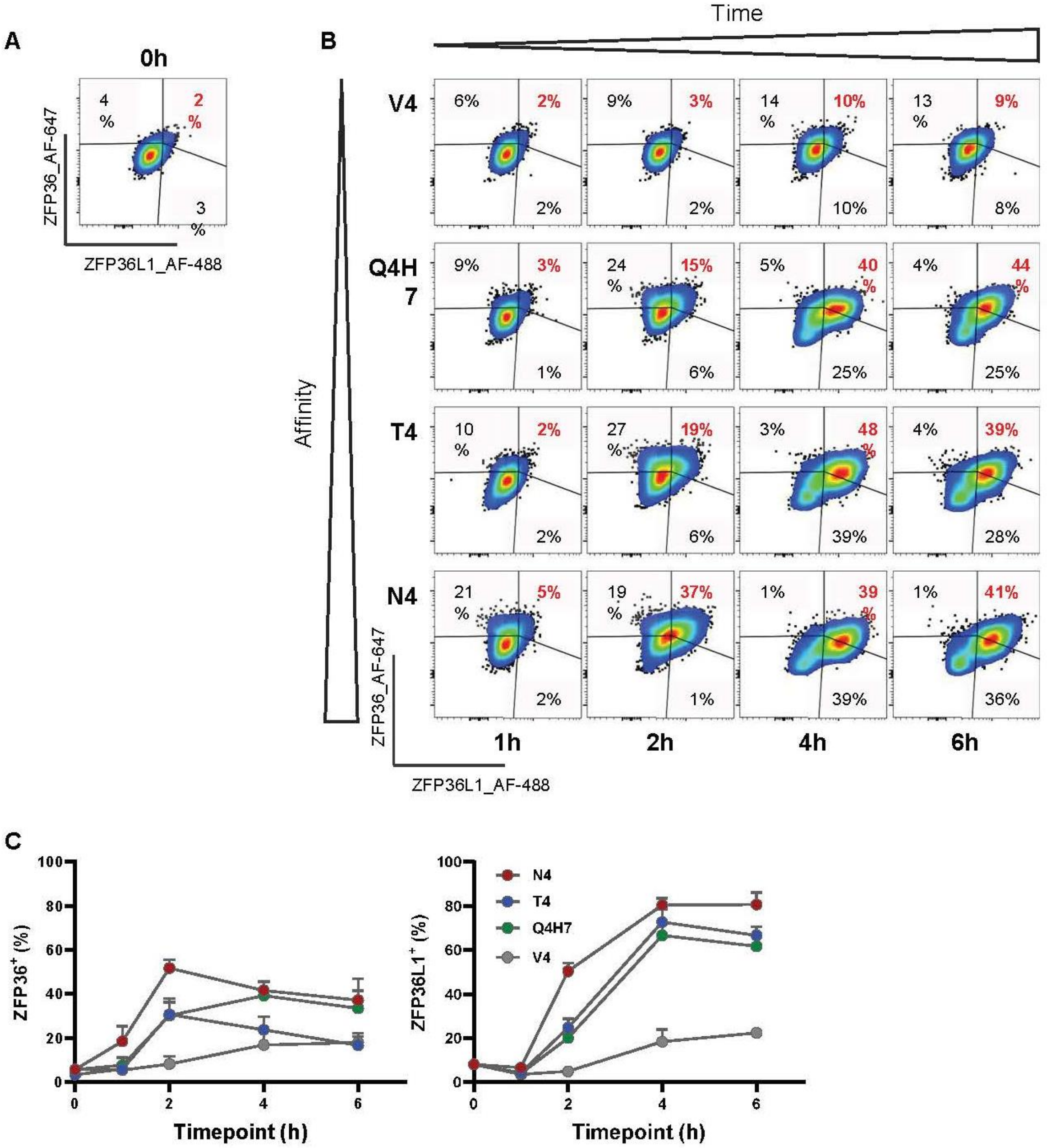


**Figure S1. ZFP36 and ZFP36L1 expression exhibit distinct kinetics and sensitivity to antigen affinity.** Representative flow cytometry plots showing staining with antibodies against ZFP36 and ZFP36L1 in one of three WT biological replicates of memory-like T cells when (**A**) unstimulated, or (**B**) after stimulation for up to six hours with 0.1 nM N4, T4, Q4H7, or V4 . (**C**) The frequencies of ZFP36- and ZFP36L1-expressing cells were plotted; error bars show standard deviation of the mean (SD) of three biological replicates.

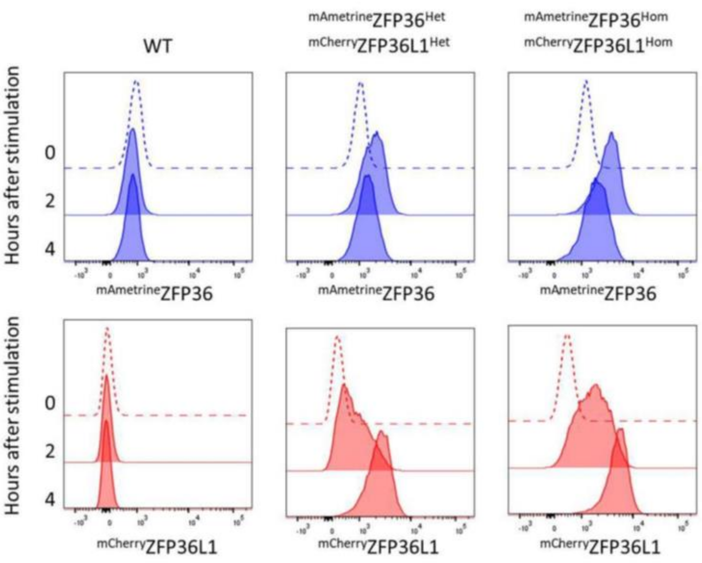


**Figure S2. Homozygous memory T cells are optimal for detection of $^{\text{mAmetrine}}$ZFP36 and $^{\text{mCherry}}$ZFP36L1 expression.** Wildtype (WT), $^{\text{mAmetrine}}$ZFP36$^{\text{Het}}$ $^{\text{mCherry}}$ZFP36L1$^{\text{Het}}$ or $^{\text{mAmetrine}}$ZFP36$^{\text{Hom}}$ $^{\text{mCherry}}$ZFP36L1$^{\text{Hom}}$ memory-like T cells were stimulated with 0.1 nM N4 for up to four hours for analysis by flow cytometry. Memory-like cells from one biological replicate per genotype was analysed.

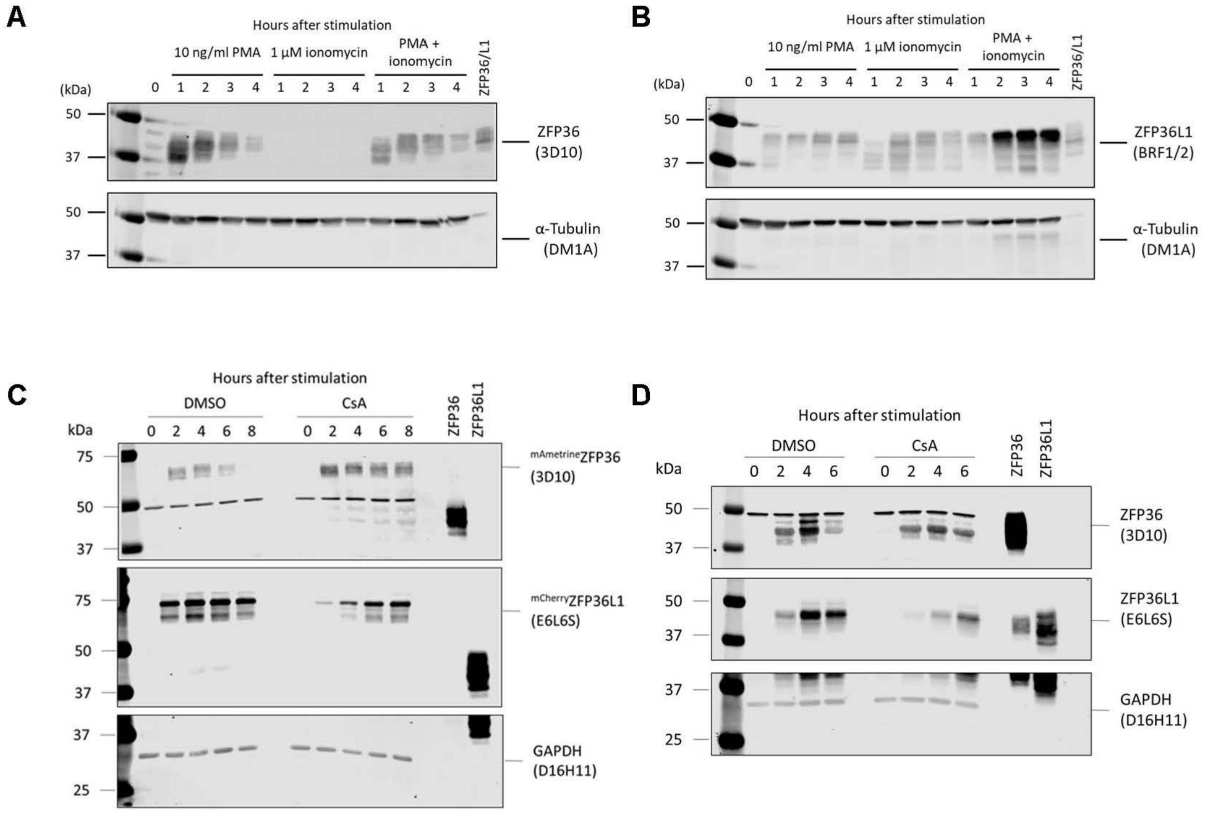


**Figure S3. Western blot analysis corroborates flow cytometry data that CsA increases ZFP36 and inhibits ZFP36L1 expression.** (**A - B**) Wildtype (WT) OT1 memory-like T cells were stimulated for up to four hours with 10 ng/ml PMA and/or 1 µM ionomycin or cultured with DMSO as a control. 30 µg of protein was loaded per lane. (**C**) $^{mAmetrine}$ZFP36$^{Hom}$ $^{mCherry}$ZFP36L1$^{Hom}$ or (**D**) WT OT1 memory-like T cells were pre-treated with 500 ng/ml CsA or DMSO control before being stimulated with 0.1 nM N4 ± CsA for up to 8 hours. (**C**) 14 µg or (**D**) 20 µg of protein was loaded per lane. (**A - D**) Immunoblots were analysed using anti-ZFP36 (3D10), anti-ZFP36L1 (BRF1/2) or anti-ZFP36L1 (E6L6S), anti-α-tubulin (DM1A) or anti-GAPDH (D16H11) antibodies as indicated. Lysates from HEK 293T expressing FLAG-tagged murine ZFP36 or ZFP36L1 were used as positive controls for blotting where 1 µg of protein was loaded per lane.

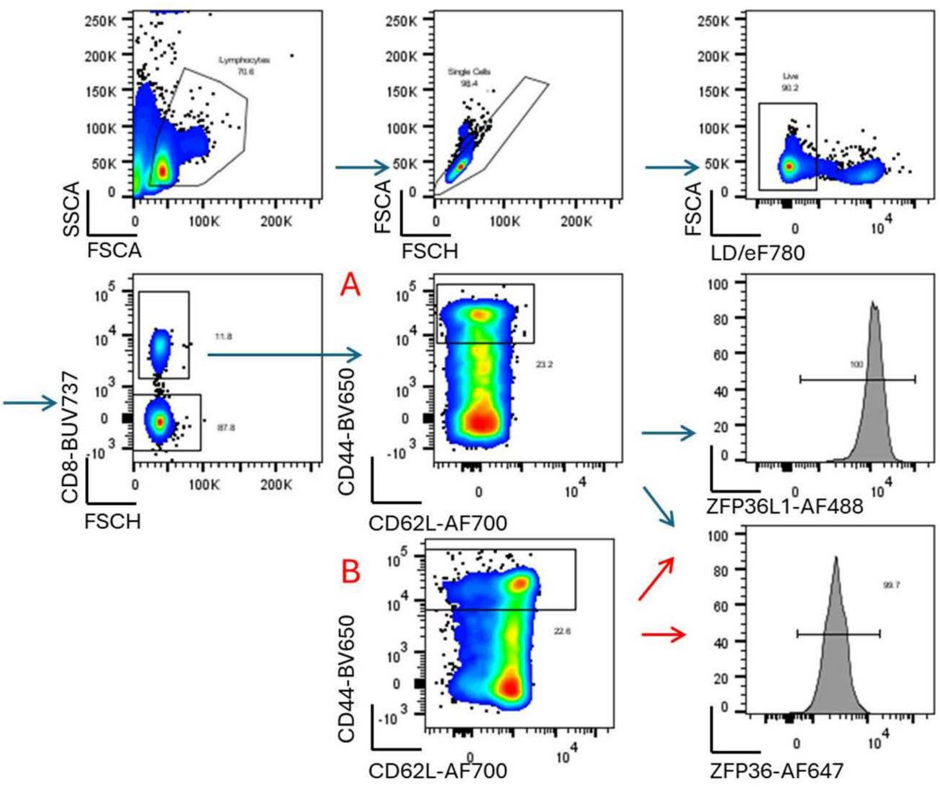


**Figure S4. *Ex vivo* memory T cell flow cytometry gating strategy.** Splenocytes from wildtype (WT) mice were gated for analysis of the memory T cell population as follows. The 'lymphocytes' and 'Single cells' gate was used to exclude debris and doublets respectively. Live cells were gated according to lack of staining with the viability dye eFluor780. Live CD8$^{+}$ T cells were then gated for CD44$^{high}$ CD62L$^{low}$ expression which are markers on memory T cells. ZFP36 and ZFP36L1 expression was analysed in memory T cells (**A**) stimulated for three hours with 10 ng/ml PMA and 1 µM ionomycin and (**B**) unstimulated cells.

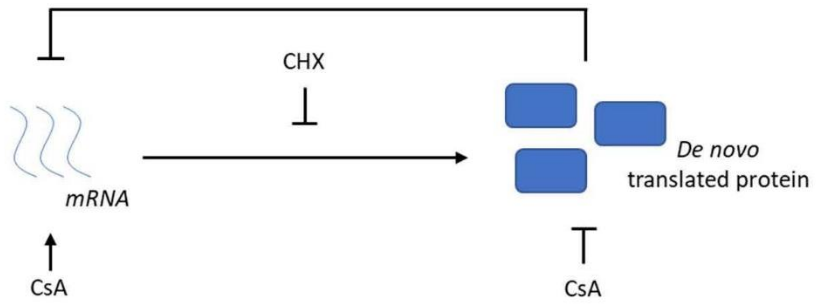


**Figure S5. Cycloheximide increases the abundance of mRNA which is negatively regulated by *de novo* translated proteins.** For mRNA transcripts that are subject to negative regulation by *de novo* translated proteins, inhibition of translation by cycloheximide results in increased mRNA abundance due to reduced suppression. Regulation of *Zfp36* mRNA by cyclosporin A (CsA) may be the result of increasing the activity of *de novo* translated proteins which suppress *Zfp36* mRNA expression. Alternatively, CsA may act independently of *de novo* protein translation. For example, CsA may promote the transcription of *Zfp36*.

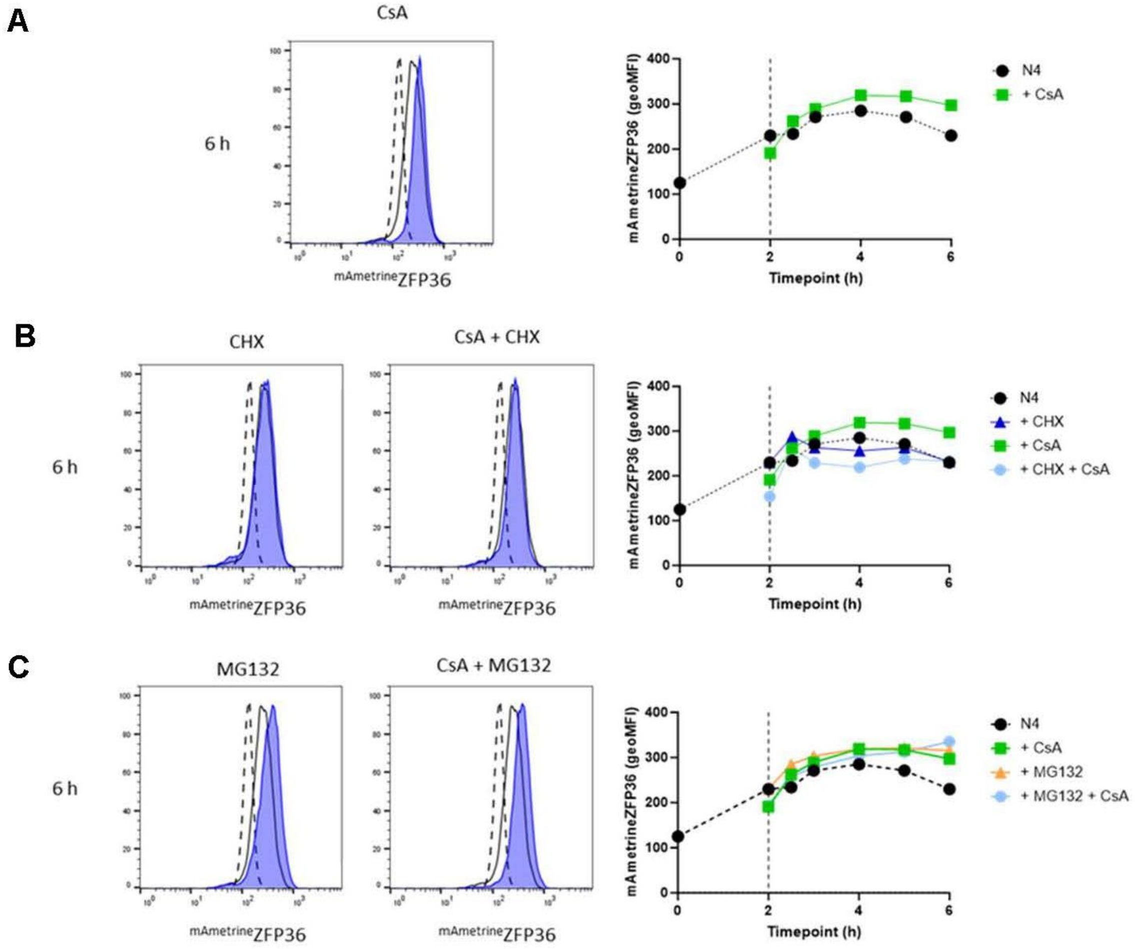


**Figure S6. Increased ZFP36 protein abundance in the presence of CsA is not due to differences in protein stability or degradation.** Memory-like T cells were pre-treated for 30 minutes with 500 ng/ml CsA or DMSO alone before being stimulated with 0.1 nM N4 ± CsA for up to 6 hours. After two hours of stimulation ± CsA, (**B**) 10 µg/ml CHX (translation inhibitor) or (**C**) 10 µM MG-132 (proteasome inhibitor) was added and $^{mAmetrine}$ZFP36 expression measured 30, 60, 120, 180 and 240 minutes after the addition of the CHX or MG-132. Graphed data shows $^{mAmetrine}$ZFP36 geoMFI of live cells from one homozygous biological replicate.
